# An assembly of genomic dataset sequences of the sugar beet root maggot *Tetanops myopaeformis*, TpSBRM_v1.0

**DOI:** 10.1101/2023.10.11.561894

**Authors:** Nadim W. Alkharouf, Chenggen Chu, Vincent P. Klink

**Affiliations:** Department of Computer and Information Sciences, Towson University, Towson, MD, 21252, USA; USDA-ARS-NA- Northern Great Plains Research Laboratory, 1307 N 18TH ST Northern Crop Science Laboratory, Fargo, ND 58102, USA; USDA-ARS-NEA-BARC, Molecular Plant Pathology Laboratory, Building 004, Room 122, BARC-West, 10300 Baltimore Ave., Beltsville, MD 20705, USA

**Keywords:** Draft genome, *Tetanops myopaeformis*, sugar beet root maggot, sequences, *Beta vulgaris*, sugar beet

## Abstract

The sugar beet root maggot (SBRM), *Tetanops myopaeformis* (von Röder), is a devastating pathogen of sugar beet (SB), *Beta vulgaris*, ssp vulgaris (*B. vulgaris*), an important food crop, while also being one of only two plants globally from which sugar is widely produced, and accounting for 35% of global raw sugar with an annual farm value of $3 billion in the United States alone. SBRM is the most devastating pathogen of sugar beet in North America. The limited natural resistance of *B. vulgaris* necessitates an understanding of the SBRM genome to facilitate generating knowledge of its basic biology, including the interaction between the pathogen and its host(s). Presented is the de novo assembled draft genome sequence of *T. myopaeformis* isolated from field-grown *B. vulgaris* in North Dakota, USA. The SBRM genome sequence will also be valuable for molecular genetic marker development to facilitate host resistance gene identification and knowledge, including SB polygalacturonase inhibiting protein (PGIP), and development of new control strategies for this pathogen.

**SPECIFICATIONS:** Organism/cell line/tissue: *Tetanops myopaeformis*

Sequencer or array type: PacBio Revio flow cell

Data format: Raw and processed

Experimental factors: DNA extracted from a wild-type strain, no treatment

Experimental features: Genome sequencing

Consent n/a

Sample source location: Sugar beet field at Fargo, North Dakota, USA

**DIRECT LINK TO DEPOSITED DATA:** BioSample accession: SAMN37733483

Temporary Submission ID: SUB13882507

BioProject ID: PRJNA1026092

Release date: 2024-09-09, or with the release of linked data, whichever is first

## INTRODUCTION

*Tetanops myopaeformis* (von Röder), the sugar beet root maggot (SBRM), is a devastating pathogen of sugar beet (SB), *Beta vulgaris*, ssp vulgaris (B. vulgaris) (Hein et al. 2009; Fugate et al. 2019). SB is an important food crop, while also being one of only two plants globally from which sugar is widely produced, and accounting for 35% of global raw sugar with an annual farm value of $3 billion in the United States alone (Hein et al. 2009; Fugate et al. 2019). SBRM is the most devastating pathogen of sugar beet in North America (Hein et al. 2009; Fugate et al. 2019). Agricultural control of SBRM is limited by a scarcity of genetic knowledge. The analysis presented here is aimed at providing such a resource for the scientific community and a basis for future updates. The work describes the data, explains its utility to the community, provides protocols and references, and provides a documented link to the data that is in a standard, re-useable format.

## MATERIALS and METHODS

### DNA isolation and sequencing

DNA isolation and sequencing of the agriculturally important *T. myopaeformis* is performed at CD Genomics using their proprietary methods. Briefly, DNA is isolated from liquid nitrogen flash frozen larvae. For sample preparation and DNA quality control, isolated DNA quality and quantity are assessed using Qubit and Agilent 5200 Fragment Analyzer. For DNA fragmentation, the DNA is cut into 15 Kb or larger fragments using the Covanis g-TUBE, and subsequently purified using magnetic beads. For DNA repair and end modification, DNA repair enzyme is used to correct any DNA damage and ensure uniformity. Furthermore, the overhang adapters are ligated to the end of the repaired DNA ends, followed by purification using magnetic beads. For SMRTbell library preparation, the repaired and adapter-ligated DNA fragments are then converted into SMRTbell libraries. For SMRTbell library size selection, the SMRTbell libraries are subjected to BluePippin size selection to enrich the fragments over 9-13 Kb and 15 Kb. For binding to polymerase, the size-selected SMRTbell libraries are bound to DNA polymerase molecules. For DNA sequencing, the prepared SMRTbell libraries, with bound polymerase molecules, are loaded onto the PacBio Sequel Revio sequencing platform with options enabled to retain subreads and kinetics sequencing metrics. Furthermore, BAM files generated through HiFi sequencing are converted to FastQ files using the SamTools Fastq algorithm. The FastQ files are imported for genome assembly.

### Assembly of genomic fragments

A draft of the *T. myopaeformis* genome is assembled, BioSample accession: SAMN37733483, Temporary SubmissionID: SUB13882507, BioProject ID PRJNA1026092. The mate-pair library produces a total number of raw reads of 6,356,906. The total read length is 71,844,227,661 and the N50/N90 reads are 11313 and 8294, respectfully. The assembly statistics show a total length of 414,327,873 with the number of contigs of 8,228. The contigs N50 is 57,402. The largest contig is 573,329 bp. The mean genome coverage is 94x.

## DATA ANALYSIS and RESULTS

### Assembly of *T. myopaeformis* genomic sequences

The *T. myopaeformis* sequencing and assembly statistics are summarized (Table 1). PacBio HiFi reads were assembled using the pipeline Flye, version 2.9.2, described in Kolmogorov et al. (2019). Default values were used, except for setting the --asm-coverage argument to 50, to reduce memory consumption. Flye was installed and run on the Windows Subsystem for Linux (Ubuntu 22.04) running on a Windows 2022 workstation with 45 GB of memory.

**Table 1.** *T. myopaeformis* TpSBRM_v1.0 genome assembly statistics.

|  |  |
| --- | --- |
| Sequencing technology: | PacBio HiFi |
| Total number of raw reads: | 6,356,906 |
| Total read length: | 71,844,227,661 |
| Reads N50/N90: | 11,313 / 8,294 |
| GC% | 40.2 |
| **Assembly statistics:** |  |
| Total length: | 414,327,873 |
| Number of contigs: | 8,228 |
| Contigs N50: | 57,402 |
| Largest contig: | 573,329 |
| Mean coverage: | 94 |

## ACKNOWLEDGEMENTS

This work is supported by the USDA-ARS NP 8042-21220-262-000D project to VK. The mention of trade names or commercial products in this publication is solely for the purpose of providing specific information and does not imply recommendation or endorsement by the United States Department of Agriculture. USDA is an equal opportunity provider and employer.

